# Olfactory marker protein regulates refinement of the olfactory glomerular map

**DOI:** 10.1101/309401

**Authors:** Dinu F. Albeanu, Allison C. Provost, Prateek Agarwal, Edward Soucy, Venkatesh N. Murthy

**Affiliations:** Cold Spring Harbor Laboratory, Cold Spring Harbor, NY 11724, USA; Department of Molecular & Cellular Biology, Harvard University, Cambridge, MA, USA; Center for Brain Science, Harvard University, Cambridge, Massachusetts, USA

**Author notes:** These authors contributed equally (Dinu F. Albeanu, Allison C. Provost, Prateek Agarwal). **Corresponding Author**: Venkatesh N. Murthy, Harvard University, Department of Molecular & Cellular Biology, 16 Divinity Ave., Cambridge, MA 02138.

## Abstract

The olfactory glomerulus is the anatomical and functional unit of the olfactory bulb, defined by convergent input from olfactory sensory neuron (OSN) axons expressing the same type of odorant receptor (OR). A key marker of mature OSNs is the olfactory marker protein (OMP), whose deletion has been associated with deficits in OSN signal transduction and odor discrimination. Here, we have investigated glomerular odor responses and anatomical architecture in mice in which one or both alleles of OMP were replaced by the fluorescent synaptic activity reporter, synaptopHluorin (OMP^+/−^ and OMP^−/−^ mice, respectively). Functionally heterogeneous glomeruli, that is, ones with micro-domains with distinct odor responses were rare in OMP^+/−^ mice, but occurred frequently in OMP^−/−^ mice. Genetic targeting of single ORs revealed that these micro-domains arise from anomalous co-innervation of individual glomeruli by OSNs expressing different ORs. The glomerular mistargeting of OSNs in the absence of OMP is restricted to a local neighborhood of a few glomerular diameters. Our studies document functional heterogeneity in sensory input within individual glomeruli and uncover its anatomical correlate, revealing an unexpected role for OMP in the formation and refinement of the glomerular olfactory map.

## Introduction

OSNs reside in the olfactory epithelium where they bind odorants^1^ and send projections to the OB where axons expressing the same OR converge to form a single glomerulus on the OB surface^2–4^. This one receptor-one glomerulus rule, implies that the odor sensitivities of OSNs throughout a given glomerulus are homogeneous, defined by the OR’s odor binding properties. This overarching rule of the olfactory bulb (OB) provides the foundation for a stereotyped map across animals^5–7^.

Glomeruli, which tile the OB, are arranged in a stereotyped pattern^2,6–8^, and the two hemifields form mirror images of each other^6,8–11^ such that each OR-defined glomerulus will appear twice on the surface of each hemisphere of the OB. OSNs synapse onto principal (mitral and tufted) neurons and interneurons in the OB^11,12^. Principal neurons of the OB receive excitatory input from a single glomerulus and convey output signals that are informed by the odor sensitivities of this parent glomerulus, but also influenced by indirect inhibitory inputs from other glomeruli, top-down feedback and neuromodulatory signals^11,12^.

Intra-glomerular heterogeneity may arise from mixed OSN input to individual glomeruli. Heterogeneous glomeruli are normally rare^7,13^, but molecular or activity perturbations to olfactory sensory neurons can increase the frequency of anatomically mixed glomeruli^14–16^. Each OB hemisphere in the mouse is thought to have around 3,500 glomeruli^17^, which would be more than sufficient to accommodate two glomeruli per OR type, one per hemifield, given ~1,100 OR types. Several identified OR-specific OSNs appear to converge onto more than one glomerulus per hemifield^7,18–21^, which skews the ratio of OR number to glomerular number and could lead to heterogeneous OR glomeruli in wildtype mice.

All mature OSNs in vertebrates express a protein called olfactory marker protein (OMP), encoded by a small intronless gene^22–25^. OMP is implicated in olfactory signal transduction^26^, with a role in determining the latency and duration of odor responses by OSNs^27^. OMP has also been proposed to aid the maturation of OSNs, for example speeding up the restriction of OR genes expression to a single type^26^. A functional role for OMP was also suggested by a recent study in which glomeruli of OMP null mice respond to a wider range of odorants than expected^28^. In addition, OSNs of OMP null mice do not always terminate in the glomerular layer, but sometimes project deeper into the external plexiform layer^29^. The exact molecular function of OMP remains elusive, but pharmacological studies indicate that it functions upstream of cAMP production in OSN cilia, possibly, as a phospho-diesterase inhibitor^30^. Combined with OMP’s presence in the axon terminals of OSNs^31,32^ and the regulatory action of cAMP on OSN axon guidance^33,34^, this observation raises the possibility that OMP may regulate axon path finding and glomerular map formation.

We set out to probe the functional and anatomical glomerular heterogeneity in OMP-synaptopHluorin (spH) mice^35^, in which spH is knocked into the olfactory marker protein (OMP) locus in the genome, replacing the coding sequence of OMP. This renders OMP-spH homozygous mice (OMP^−/−^) knockouts for OMP, whereas OMP-spH heterozygotes (OMP^+/−^ mice) express one copy of the gene. Using functional imaging, as well as anatomical reconstructions, we found that OMP is necessary to preserve the one receptor-one glomerulus rule in the adult olfactory bulb.

## Results

### Absence of OMP alters odor responses, but preserves macro-organization of the OB

It has recently been shown that glomerular responses in the OMP null mice exhibit wider tuning curves than in OMP heterozygotes^36^, but these experiments utilized a very limited array of odors. This led us to ask, with a diverse odor panel, how glomerular odor representations are changed in OMP^−/−^ mice. We explored this question in OMP-SpH mice that have SpH, a synaptic activity reporter, knocked into the *OMP* locus and expressed in all OSNs. Using widefield imaging of spH signals on the dorsal surface of the bulb to a panel of 98 odorants (Suppl. Table 1), we found that single glomeruli, as determined by their spherical macrostructure (Fig. 1a), in the OMP^−/−^ mouse responded to more odors when compared to OMP^+/−^ mice (8.5 ± 0.5 in OMP^+/−^ and 9.7 ± 0.6 in OMP^−/−^; p<0.005; Fig. 1c,d, *left*). Also, the number of glomeruli responding to single odors in the panel nearly doubled in OMP^−/−^ compared to OMP^+/−^ mice (8.2 ± 0.5 in OMP^+/−^ and 16.2 ± 0.6 in OMP^−/−^; p<0.01; Fig. 1c,d, center). These differences in response patterns are indicative of pronounced changes in glomerular activity patterns in the OMP^−/−^ genetic background.

**Figure. 1.**
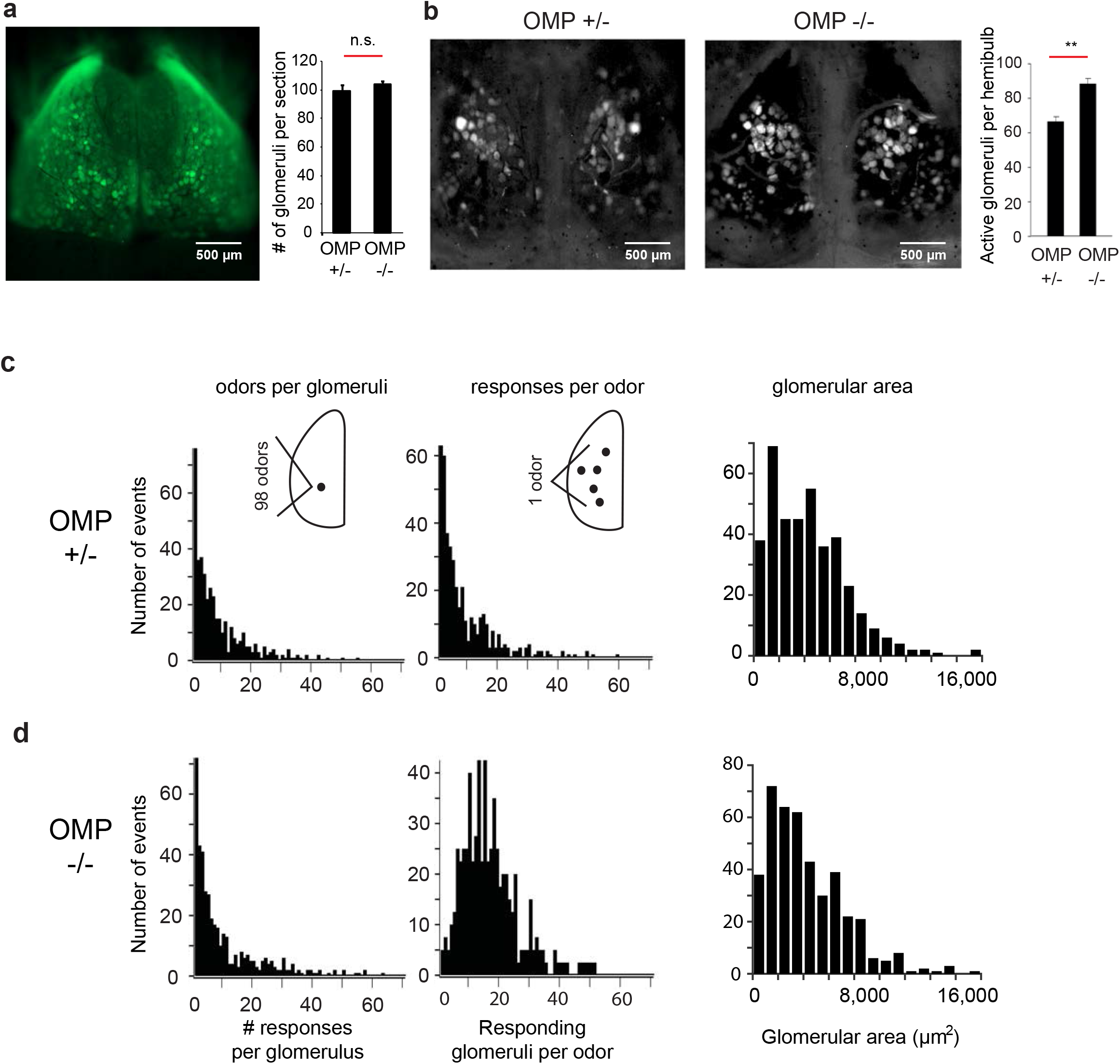
Implications for olfactory coding. **a)** (*Left*) Widefield view of dorsal surface of an OMP^+/−^ mouse shows labeled glomeruli from the resting fluorescence of spH. (*Right*) Quantification of the number of glomeruli per coronal section in central sections of the OB. 99 ± 3.44 glomeruli per coronal section for OMP+/− mice (Mean ± SEM, n = 5 slices from 2 mice), 104 ± 1.8 glomeruli per coronal section for OMP nulls (Mean ± SEM, n = 6 slices from 2 mice). **b)** (*Left*) Maximum projection glomerular odor response maps for exemplar OMP^+/−^ and OMP^−/−^ mice to a panel of 98 odorants. (Right) Average number of responsive glomeruli per hemibulb to the odor panel (Supplementary Table 1). **c,d)** (*Left*) Histograms of the number of odors to which a single glomerulus responds. (*Center*) Histograms of the number of glomeruli activated by an individual odor in the panel. (*Right*) Histograms of glomerular area in central coronal sections of the OB in OMP^+/−^ and OMP^−/−^ mice.

**Table 1.**
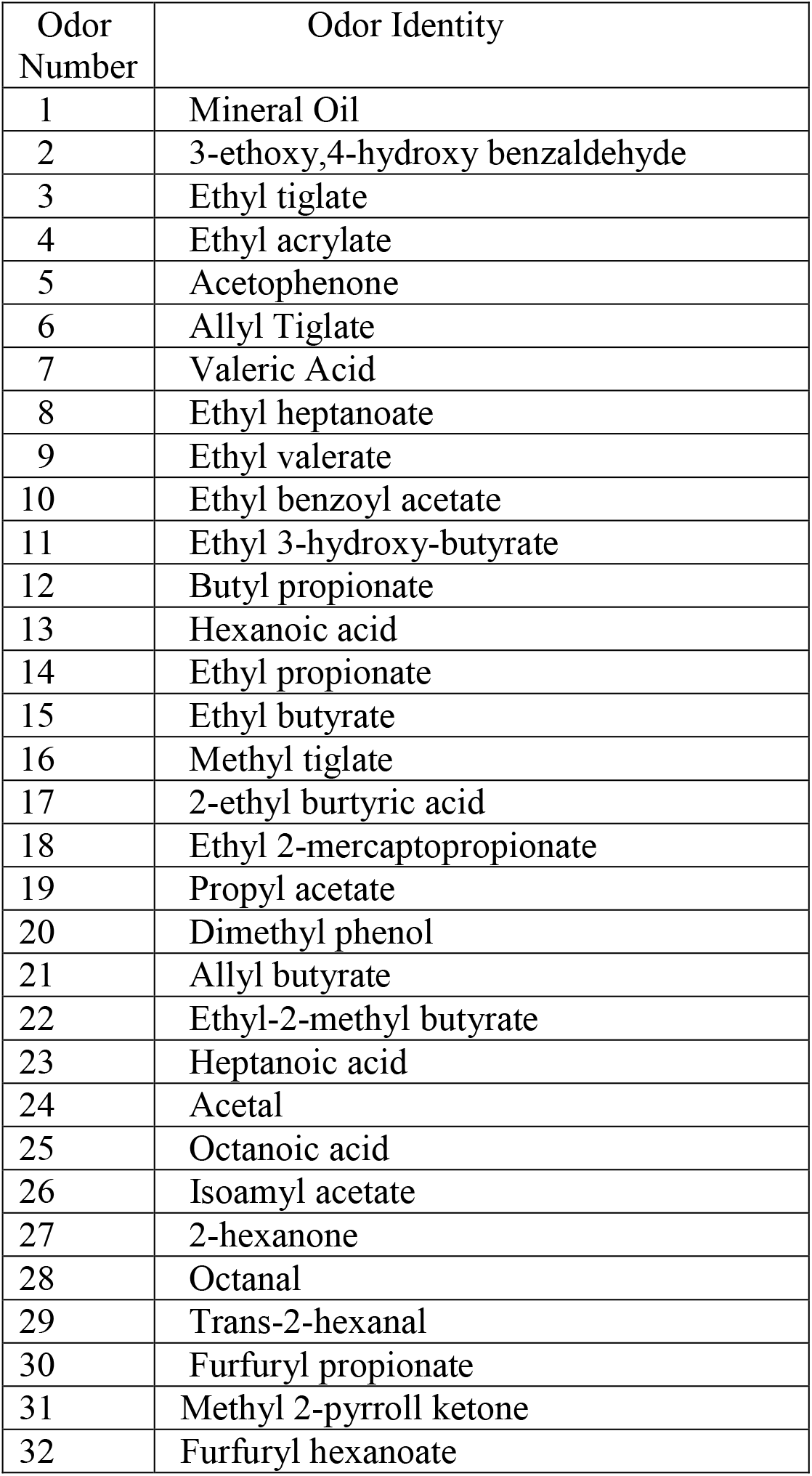
*Panel of odorants used for the two-photon experiments*.

| Odor Number | Odor Identity |
| --- | --- |
| 1 | Mineral Oil |
| 2 | 3-ethoxy,4-hydroxy benzaldehyde |
| 3 | Ethyl tiglate |
| 4 | Ethyl acrylate |
| 5 | Acetophenone |
| 6 | Allyl Tiglate |
| 7 | Valeric Acid |
| 8 | Ethyl heptanoate |
| 9 | Ethyl valerate |
| 10 | Ethyl benzoyl acetate |
| 11 | Ethyl 3-hydroxy-butyrate |
| 12 | Butyl propionate |
| 13 | Hexanoic acid |
| 14 | Ethyl propionate |
| 15 | Ethyl butyrate |
| 16 | Methyl tiglate |
| 17 | 2-ethyl burtyric acid |
| 18 | Ethyl 2-mercaptopropionate |
| 19 | Propyl acetate |
| 20 | Dimethyl phenol |
| 21 | Allyl butyrate |
| 22 | Ethyl-2-methyl butyrate |
| 23 | Heptanoic acid |
| 24 | Acetal |
| 25 | Octanoic acid |
| 26 | Isoamyl acetate |
| 27 | 2-hexanone |
| 28 | Octanal |
| 29 | Trans-2-hexanal |
| 30 | Furfuryl propionate |
| 31 | Methyl 2-pyrroll ketone |
| 32 | Furfuryl hexanoate |

In order to better understand the functional changes in glomerular responses, we also examined whether the loss of OMP leads to changes in the macro-organization of glomeruli.

Does the increased number of responding glomeruli indicate the presence of more glomeruli in OMP^−/−^ mice? Using fixed tissue slices, we counted the number of glomeruli present in the most central OB slices. Glomerular borders were clearly indicated by DAPI staining of the vast number of juxtaglomerular cell nuclei surrounding the glomerulus^37^. We found that the total number of glomeruli did not significantly change between OMP^+/−^ and OMP^−/−^ mice (99 ± 3.4 and 104 ± 1.8 glomeruli per OB slice, respectively, Fig. 1a, left; p>0.65, Kolmogorov-Smirnov test). Moreover, the size of glomeruli in the OMP^+/−^ and OMP^−/−^ mice was very similar (4,303 ± 149 μm^2^ and 4,266 ± 149 μm^2^, respectively Fig. 1c,d, *right*; p>0.3, Kolmogorov-Smirnov test), corresponding to glomerular average diameters of 74.0 μm and 73.7 μm, consistent with previous studies^38,39^. Together, our results indicate that the macro-structure of the glomerulus is relatively unperturbed in the absence of OMP. The increased promiscuity of glomerular odor responses, but lack of change in the anatomical macro-organization of glomeruli, led us to hypothesize that OMP plays a role in shaping the organization of OSN axons within individual glomeruli.

### Functional microdomains in OMP null mice

We asked how responses within single glomeruli change in the OMP null background. We used two-photon laser scanning microscopy (2PLSM) to examine the microstructure of glomerular responses to odors in OMP-spH mice^35,40^. Glomerular activity in each field of view was probed serially with a panel of 32 odorants (Suppl. Table 1). A spatially contiguous region of resting fluorescence surrounded by dark voids was taken to be an individual glomerulus (Fig. 2a).

**Figure. 2.**
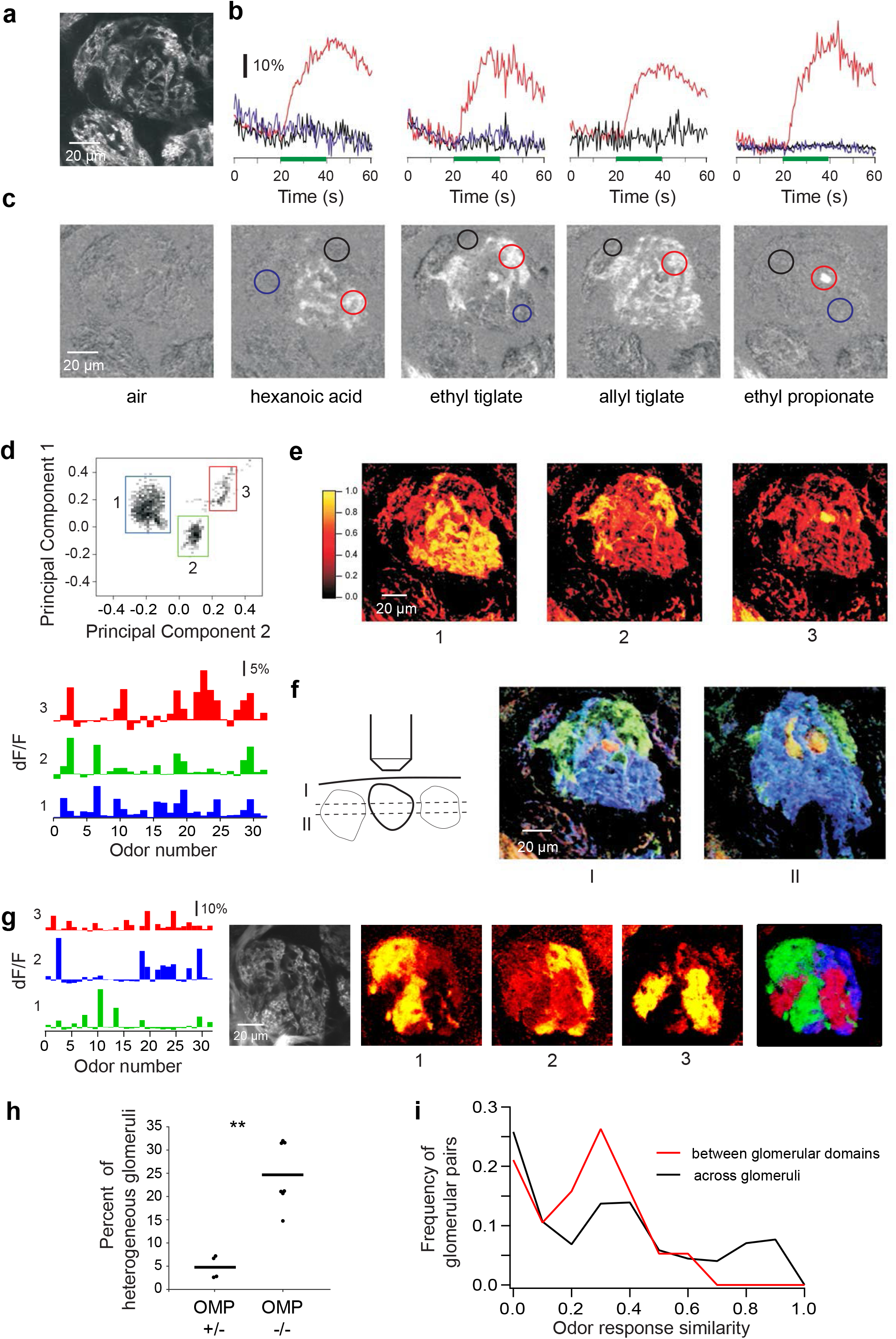
Analysis of functional glomerular heterogeneity in OMP^−/−^ glomeruli. **a)** Raw two photon fluorescence of a glomerulus in an OMP^−/−^ mouse. **b, c)** Example time courses and average responses (dF/F) of the glomerulus in (a) to 5 different stimuli. Colored circles mark regions of interest. Horizontal line marks stimulus delivery. **d)** (*Top*) Projection of all pixels in **(a)** on two principal components (PC1 vs. PC2) reference coordinates. PCA was performed on the odor response spectra of each pixel. (*Bottom*) Average odor response spectra of three functional clustered identified in **(a)**. **e)** Spatial correlograms showing the spatial distribution of the three pixel clusters identified via PCA **(d)** within the example glomerulus **(a)** were obtained by correlating the average response vectors of the identified clusters to the odor responses of individual pixels in the field of view. Numbers mark correlation maps corresponding to each functional cluster in **(d)**. Note that each cluster corresponds to a spatially contiguous area we refer to as a microdomain. **f)** (*Left*) Cartoon depicting two optical imaging planes (*z*) within the example glomerulus (a) at two different depths (I & II, 20 μm apart). (*Center & Right*) RGB color scheme overlays of the correlograms determined as in **(e)** for the two optical planes sampled within the glomerulus. **g)** Three spatial microdomains in a second example glomerulus in the OMP^−/−^ mouse. (*Left*) Average odor response spectra of functional clusters identified via PCA. (*Right*) Resting fluorescence, single pixel correlograms, and their overlay in an RGB scheme. **h)** Summary graph of the average frequency of glomerular heterogeneity recorded in OMP^+/−^ and OMP^−/−^ mice (4.8 ± 2.5%, n=4 mice versus 24.7 ± 6.9%, n=7 mice, p=0.0058, Mann-Whitney U Test; horizontal bars represent the mean, dots individual animals). **i)** Histogram of pairwise odor response similarity between functional domains within the same anatomical glomerulus (red) and between different glomeruli (black) in OMP^−/−^ mice.

Odors evoked robust increases in fluorescence intensity as described before via widefield microscopy^6,28,35,36^, as well using 2PLSM^13,35,41^. When we examined odor responses in OMP−/− mice (Fig. 2a), we frequently observed fluorescence increases within sub-regions of individual glomeruli (Fig. 2b,c). At higher magnification, we noticed that different odors activated distinct parts of a single glomerulus (Fig. 2c, Movies 1,2). Analysis of the time course of responses in different sub-regions confirmed that different odors differentially activate at least three regions within the example glomerulus shown in Figure 2b.

To obtain an objective view of the response heterogeneity within the imaged glomeruli (Fig. 2a), we used principal component analysis (PCA) to classify all the pixels in the imaged area according to their similarity. If a glomerulus is entirely homogeneous in its responses to odors, all the pixels are expected to be similar to each other. On the other hand, if different groups of pixels evoke distinct response profiles to odors, they will be separated out by PCA, even if they are not spatially contiguous. Here, each pixel was assigned an integrated odor response value (average dF/F) for each of the 32 odors, so each pixel was associated with an odor response vector.

Pixels above the signal threshold were projected on the plane determined by two of the first three principal components and it became apparent that they defined three functional clusters (Fig. 2d). Interestingly, the three functional clusters mapped onto three contiguous spatial microdomains within the glomerulus and matched the spatial domains activated by individual odorants (Fig 2e). This correspondence was revealed by correlating the average odor response vectors of the three PCA-identified functional clusters (Fig 2d, *bottom*) to the odor responses of each pixel in the field of view above the noise threshold (see Methods). Superimposing the three PCA-identified cluster of pixels (using an RGB color-code) fully reconstructed the glomerular anatomy assessed from the resting fluorescence image (Fig. 2f), leaving no empty regions.

To understand the three-dimensional structure of this heterogeneous glomerulus, we performed PCA again on single pixel responses for a different optical plane (20 μm more superficial) within the same glomerulus (Fig. 2f). The same three functional domains were apparent in this second optical plane, only at different relative spatial proportions. In the superficial optical plane, the ‘blue’ microdomain dominated the glomerulus response, pushing aside ‘green’ response territory. Similar exemplar data are shown for several other glomeruli in Fig. 2g and Supplementary Fig.1 (also see Movies 3,4). The spatial structure of the functional microdomains identified suggest that these clusters are due to different types of OSN inputs innervating the same glomerulus. Another argument in favor of multi-OSN type glomerular innervation is that in many instances, such as the z-stack shown in Supplementary Figure 2 (also Movies 5-7), we could follow different bundles of OSN fibers of slightly different resting fluorescence levels converge within the glomerulus (sampling every 2μm along the z-axis), and further relate them functionally to glomerular microdomains by eliciting differential responses with the panel of odorants.

To determine the frequency of occurrence of mixed glomeruli, we outlined anatomical boundaries of glomeruli using the resting fluorescence images. We then counted the total number of active glomeruli in the imaged region (Methods). Functionally heterogeneous glomeruli were identified in both OMP^+/−^ and OMP^−/−^ animals using PCA. Functionally mixed glomeruli occurred at significantly higher rate (24.7 ± 6.9%, n=7 mice) in the OMP^−/−^ mice, compared to OMP^+/−^ mice (4.8 ± 2.5%, n=4 mice, Fig. 2h, Mann-Whitney U Test, p=0.0058). We further calculated the pairwise odor response similarity between functional domains observed within the anatomical glomeruli versus across random pairs of glomeruli (including those that are heterogeneous) in the fields of view sampled in OMP^−/−^ mice. The intra-glomerular microdomains were no more similar to each other in their odor tuning than random pairs of distinct anatomical glomeruli (Fig. 2i).

To further investigate how the apparent mixing of multiple types of OR fibers alters the glomerular map in OMP^−/−^ mice, we sampled larger fields of view tiling the surface of the bulb. We found that functionally fragmented glomeruli were generally located near glomeruli that responded uniformly to the odorant panel and were functionally matched to one micro-domain of the mixed glomeruli (Fig. 3a). In many instances, within a few glomerular lengths, we also identified additional glomerular fragments that had similar tuning to the odors in our panel (Fig. 3b and Supplementary Fig. 3). By contrast, in OMP^+/−^ mice, the occurrence of functionally heterogeneous glomeruli and of the glomerular duplicates was significantly lower (Fig. 2h, Supplementary Fig. 4)

**Figure. 3.**
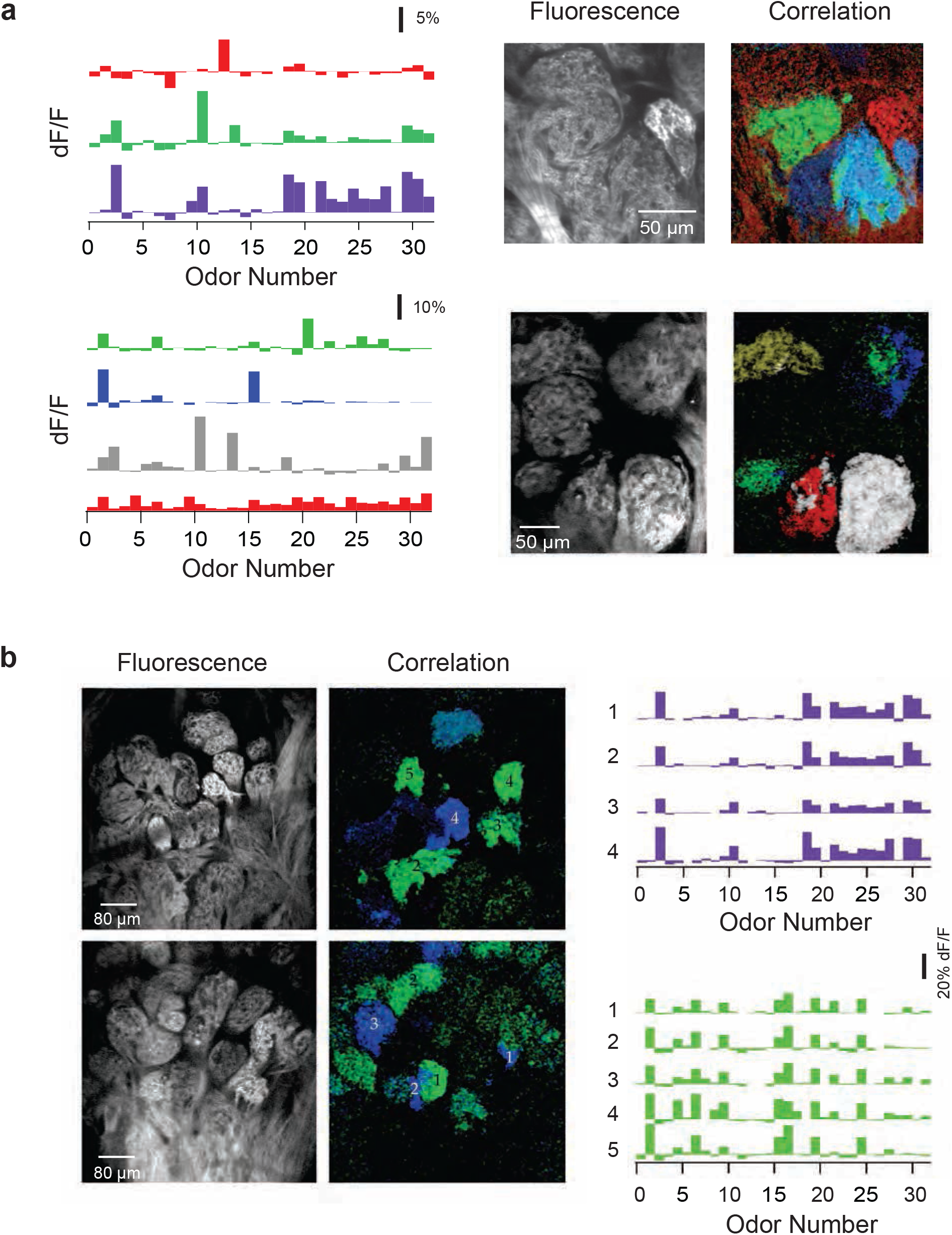
Local functional glomerular duplications in OMP^−/−^ mice. **a)** (*Left*) Example average odor response spectra of PCA-identified functional clusters corresponding to glomerular microdomains from two different fields of view (top vs. bottom). (*Right*) Corresponding resting fluorescence and overlay of correlograms. Note the presence of functionally homogeneous and heterogeneous glomeruli adjacent to each other. For example, in the lower panel, six anatomically identifiable glomeruli are discernable. The correlograms (*Right*) indicate that green microdomains are shared between three close-by anatomical glomeruli. White and red microdomains mix within one anatomical glomerulus. The white cluster is also present as a spatial-functional homogeneous adjacent glomerulus. **b)** Two partially overlapping example fields of view in one OMP^−/−^ mouse. (*Left*) Resting glomerular fluorescence; (*Center*) Correlograms: colors correspond to functionally matched glomeruli or subglomerular microdomains; (*Right*) Odor response spectra for the green and blue glomeruli. Numbers correspond to the location of the functionally similar glomeruli in the sampled fields of view.

### Anatomical microdomains are prominent in OMP^−/−^ mice

We hypothesized that the functional microdomains observed in OMP-null mice were due to anatomical heterogeneity. Specifically, we predicted that multiple types of OSNs, as defined by the ORs they express, converge onto a single glomerular structure To test this hypothesis, we used mouse lines with single labeled glomeruli, M72-RFP^42^ and P2-LacZ^20^, crossed into the OMP-spH background, and compared the glomerular convergence in OMP^+/-^ mice to OMP null littermates. In heterogeneous glomeruli, we expect to see genetically labeled OR-specific OSN axons intermixed with unlabeled OSN axons (if they were expressing a different, untargeted OR).

As expected, glomeruli from OMP^+/−^ mice were generally homogeneous (Fig 4a). By contrast, in OMP^−/−^ mice, partially filled, heterogeneous glomeruli were frequently identified (Fig. 4b). In order to quantify heterogeneity within a single glomerulus, we determined the overlap between P2 β-galactose signal (pseudocolored red in Fig. 4a) and OMP-spH signal (green in Fig. 2,3, see Methods). Glomerular borders were determined by the cluster of nuclei or juxtaglomerular cells, stained by DAPI (Fig. 4a). We were then able to express glomerular homogeneity as the ratio of number of spH-positive pixels that were also P2 positive and the number of total spH positive pixels. This quantifies the percentage of pixels in a glomerulus that express our target gene. P2 positive fibers frequently ramified into anatomical microdomains of the target glomerulus of OMP^−/−^ mice, on average filling 61% of the glomerulus (n=52 glomeruli) compared with 82% in OMP^+/−^ mice (n=80 glomeruli) (p<0.001, Kolmogorov Smirnov test; Fig. 4c).

**Figure. 4.**
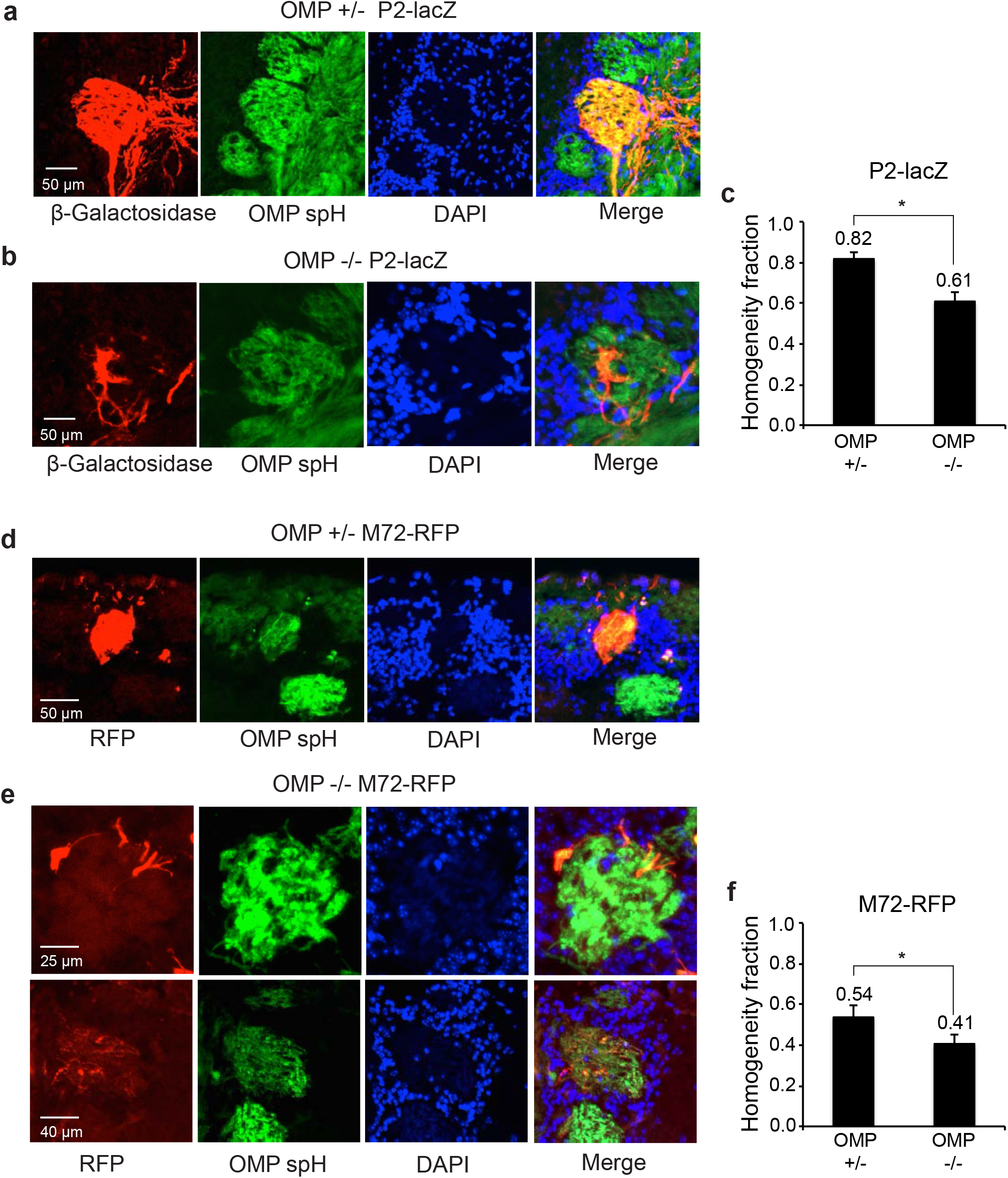
Anatomical glomerular heterogeneity in the OMP^−/−^ mouse. **a)** Example homogenous glomerulus in a P2-lacZ OMP^+/−^ mouse. P2-positive OSN axons labeled with β-Gal (red), SpH-positive OSN axons (green) and glomerular outline by DAPI-stained nuclei of juxtaglomerular cells (blue). Note the high overlap between the red and green pixels in the merged image. **b)** Example heterogeneous glomerulus in a P2-lacZ OMP^−/−^ mouse. P2-positive OSN axons labeled with β-Gal (red) do not overlap entirely with SpH-positive OSN axons (green) and only partially fill the glomerulus, outlined by DAPI-stained nuclei of juxtaglomerular cells (blue). **c)** Quantification of glomerular homogeneity (P2-lacZ) in the OMP^+/−^ versus OMP^−/−^ mice. Homogeneity fraction: 0.82 ± 0.03 (Mean ± SEM, n = 52 glomeruli) in OMP^+/−^ mice, and 0.61± 0.05 (Mean ± SEM, n = 40 glomeruli) in OMP^−/−^ animals. **d,e)** Same as a) and b) an M72-RFP example glomerulus in the OMP^+/−^ and OMP^−/−^ genetic backgrounds. **f)** Quantification of glomerular homogeneity (M72-RFP) in OMP^+/−^ versus OMP^−/−^ mice. Homogeneity fraction: 0.54 ± 0.06 (Mean ± SEM, n = 19 glomeruli) in OMP^+/−^ mice, and 0.41 ± 0.05 (Mean ± SEM, n = 18 glomeruli) in OMP^−/−^ animals.

In the M72-RFP line, we also observed that M72-RFP OMP +/− mice often exhibited filled, homogenous glomeruli (Fig. 4d). Interestingly, the M72-RFP line showed a distinct phenotype of heterogeneity, with discrete subdomains sometimes visible (Fig. 4e *top*), but often displaying wider, more diffuse borders (Fig. 4e bottom row). We did not amplify RFP signals using secondary conjugates, which led to poorer detection efficiency than for spH signals. Therefore, the percent overlap even in OMP^+/−^ mice was low, and we relied on relative measures. M72-RFP glomeruli had significantly higher homogeneity fraction in OMP^+/−^ than in OMP^+/−^ mice (54%, n=19 glomeruli versus 41%, n=19 glomeruli, p<0.05, Kolmogorov Smirnov test; Fig. 4f).

### Duplicate glomeruli are more frequent in OMP null mice

In addition to microdomains within single glomeruli, we also found functional evidence of duplicate glomeruli, found in close proximity to the target sites (Fig. 3). This duplication could arise from OSNs expressing a particular OR terminating in multiple glomeruli in the vicinity of each other. Gene targeted glomeruli are found in well-defined locations on the OB surface, and there are usually two, one per hemifield, though low frequency local duplications have been reported even in adult wild-type mice^7^. To determine the number of labeled glomeruli in each hemifield, we imaged OBs in whole mount preparation (Fig. 5a). On both the OMP^−/−^ and OMP^+/−^ background, M72-RFP glomeruli were always located on the dorsal surface, segregating to both the medial and lateral hemifields (Fig. 5a,b left panels). We never observed ectopic glomeruli that terminated well beyond the expected target region. As expected, in the OMP^+/−^ background M72-RFP glomeruli were usually limited to one per hemifield (Fig. 5a, top), but the M72-RFP glomerulus appeared frequently duplicated in OMP^−/−^ mice (Fig. 5a, bottom). Overall, glomerular duplication in the M72-RFP line occurred more frequently in OMP^−/−^ mice than OMP^−/−^ littermates, with on average of 1.36 ± 0.07 (Mean ± SEM, n = 76 hemibulbs) versus 1.05 ± 0.03 (n=56 hemibulbs) M72-RFP glomeruli per hemifield, respectively (p<0.001, Wilcoxon rank-sum test; Fig. 5b). Importantly, duplicate glomeruli were always located within a few average glomerular diameters of each other.

**Figure. 5.**
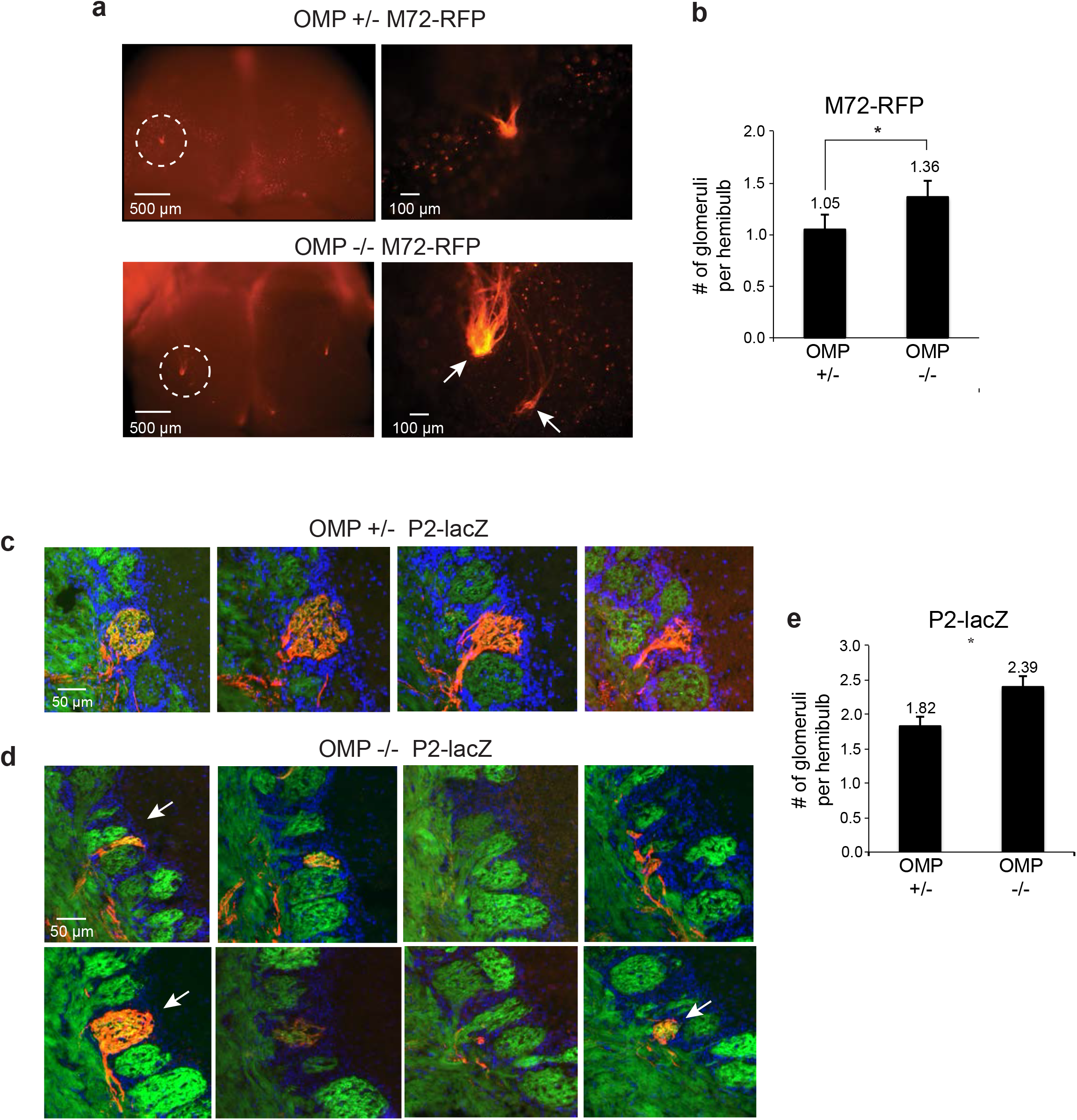
Duplicate glomeruli in the OMP null mouse. **a)** (*Left*) Whole mount of the OB dorsal surface in a M72-RFP OMP^+/−^ (Top) and OMP^−/−^ (Bottom) mouse with a the M72 glomerulus fluorescently labeled. *(Right)* Zoom-in of circled region. Note the duplicated glomeruli in the OMP^−/−^ mouse (arrows). **b)** Quantification of duplicate glomeruli in M72-RFP mice for OMP^+/−^ and OMP^−/−^ mice. 1.05 ± 0.03 glomeruli per hemifield (Mean ± SEM, n = 56 hemibulbs) in OMP^+/−^ mice, 1.36 ± 0.07 glomeruli per hemifield (Mean ± SEM, n = 76 hemibulbs) in OMP^−/−^ mice. **c)** Serial coronal sections through individual glomeruli in one hemibulb of a P2-lacZ OMP^+/−^ mouse. P2-positive OSN axons are labeled with β-Gal (red), OSN axons with SpH (green), and nuclei of juxtaglomerular cells surrounding glomeruli by DAPI (blue). **d)** Serial coronal sections of a multiple glomeruli in one hemibulb of a P2-lacZ OMP^−/−^ mouse. Arrows indicate distinct glomeruli, and fluorescent labels as in (c). **e)** Quantification of duplicate glomeruli in P2-lacZ mice for OMP^+/−^ and OMP^−/−^ mice. 1.82 ± 0.14 glomeruli per hemifield (Mean ± SEM, n = 28 hemibulbs) in OMP^+/−^ mice, 2.39 ± 0.16 glomeruli per hemifield (Mean ± SEM, n = 28 hemibulbs) in OMP^−/−^ mice.

The P2 glomerulus has been shown to form duplicates on a wild type background^20^, and these glomeruli are positioned proximally to one another, and can be difficult to discern in whole mount brains. Because of this, we quantified the number of duplicates of P2 glomeruli in both the OMP^−/−^ and OMP^+/−^ backgrounds in fixed tissue slices. We observed one or two P2 glomeruli per hemifield in the OMP^+/−^ mice (Fig. 5c), and sometimes more than two P2 glomeruli in the OMP^−/−^ mouse (Fig. 5d). In Fig. 5d, multiple glomeruli in different tissue sections contain P2 axons (red), often in microdomains. Overall in the P2 line, glomerular duplicates occurred significantly more frequently in the OMP^−/−^ than in the OMP^+/−^ background, displaying 2.39 ± 0.16 (n=28 hemibulbs) and 1.82 ± 0.14 (n=28 hemibulbs) glomeruli per hemifield, respectively (p<0.05, Wilcoxon ranked-sum test; Fig. 5e).

These results indicate that OMP null leads to heterogeneity and duplicate glomeruli anatomically. Glomerular duplication was always local, and no gross misguidance of OSN axons at far away targets was observed in the investigated genetically targeted glomerular. These data suggest that OMP plays a role in local axon refinement at the glomerulus.

## Discussion

In this study, we present evidence of functional and anatomical glomerular heterogeneity in the main olfactory bulb of mice lacking OMP. The macrostructure of glomeruli was not altered, but within single glomeruli, functional subdomains were activated by odors, and we observed numerous local functionally duplicated glomeruli. Moreover, we identified anatomical subdomains in gene targeted mice in which OSNs expressing a particular OR also co-express a fluorescent protein. OMP^−/−^ mice tended to exhibit anatomically and functionally heterogeneous glomeruli more frequently than OMP^+/−^ mice. We also observed duplicate glomeruli in close proximity of one another, but no ectopic glomeruli were found far away from the target site indicating failure of local axon pruning, rather than radical changes in axon guidance. These findings indicate that OMP is necessary for local glomerular refinement, and the lack of OMP leads to an ‘immature’ glomerular map.

### Loss of OMP leads to heterogeneous glomeruli

High resolution imaging in our study has revealed, for the first time, functional heterogeneity within an individual glomerulus. Such heterogeneity is rare in wild-type animals, akin to what we report in OMP+/− mice, potentially explaining why a previous study using multiphoton imaging in this mouse did not uncover signs of it^13^. We also note that a substantial number of odorants may be needed to observe glomerular heterogeneity, since differential activation of individual microdomains is necessary for their identification. This suggests that the frequency of heterogeneous glomeruli we report here in the OMP^−/−^ mice represents a lower bound.

A recent report noted that tuning curves in OMP^−/−^ mice are wider than in OMP^+/−^ mice^36^. We propose that the functional and anatomical phenotype presented here accounts for increased tuning widths. Similar to Kass et al.^36^, we found that with lower resolution wide field imaging, a single glomerulus responds to more odors, and strikingly, on average a single odor evokes nearly double the number of responding glomeruli in OMP^−/−^ mice. This response pattern is explained by microdomain responses within single glomeruli that are not resolvable with wide-field imaging conditions.

Our anatomical data supports the explanation that functional microdomains recorded *in vivo* are due to the convergence of axons that express different odorant receptors into a single glomerulus. We often observed partially labeled glomeruli in gene-targeted mice and infer that the remainder of these glomeruli contain axons expressing different ORs. An alternative possibility, that the same odorant receptor has widely different odor responses and segregates differently within a glomerulus, would be a surprising finding requiring a fundamental rewriting of odor coding rules. Yet another possible explanation is related to differences in concentration of the stimulus as seen by different regions of the glomerulus, perhaps due to differences in odorant sampling at the olfactory epithelium. This possibility is unlikely since different odorants should have similar patterns of differential activation, which is not seen in our data. An additional possibility is that individual OSNs express multiple receptors in the absence of OMP^26^, causing differences among OSNs innervating a single glomerulus. This also seems unlikely as the tuning widths of single microdomains in the OMP null, and whole glomerular responses in the OMP^+/−^ mice were similar, indicating that the one neuron, one receptor rule is intact in OMP^−/−^ mice.

Together, the above observations strongly support the notion that the microdomains observed are due to convergence of multiple types of OSNs, as defined by the specific ORs they express, to individual glomeruli.

### How does the loss of OMP lead to heterogeneous glomeruli?

OMP is thought to be involved in the OSN signal transduction pathway^30^, and other studies in the OMP null have revealed slower response dynamics, decreased sensitivity, and lowered specificity to odors, similar to those of perinatal OSNs^26^. Slow dynamics of response were also reported by others^36^, but Kass, et al. report that slowed activity does not lead to decreased total activity, indicating that lack of activity may not be the culprit.

Anatomically heterogeneous and duplicate glomeruli have been reported in adolescent mice^7^ indicating that glomerular refinement occurs in young mice and homogeneous glomeruli emerge in adulthood. Moreover, activity is important in refinement and maintenance of the glomerular map^7,14^. In both studies, OSNs experienced decreased level of inputs from environment throughout postnatal maturation, and, importantly, lower spontaneous activity compared to wild-type conditions. OMP’s involvement in signal transduction may lead, in principle, to a very similar phenotype during postnatal development of the odor map, since the onset and offset kinetics of OSNs are slowed down, as is the recovery from odor adaptation. This suppression of activity has multiple downstream effects, including disrupting intraglomerular maps^43^.

An interesting case of heterogeneous glomeruli is that in the monoclonal M71 mouse that expresses the M71 OR in ~95% of its OSNs^44^ This mouse displays spherical glomeruli that tile the OB surface, rather than a single massive glomerulus. They found that single glomeruli could be anatomically heterogeneous exhibiting axons from both a gene-targeted OSN type and untargeted, non-labeled OSNs. Odor responses in this mouse are observed throughout the entire glomerular field, and discrete glomerular responses are rare. We propose that differentiable activity is important for glomerular segregation. OMP’s role in the signal transduction pathway leads to slow, blurred dynamics^30,36^ that may not allow for local differentiation. This is supported by McGann and colleagues’ enrichment paradigm in which OMP-spH null odor responses become sparser after olfactory enrichment^28^. We posit that enrichment leads to increased activity that allows for more efficient differentiation between OSN types. The resulting reduction in glomerular heterogeneity leads to fewer responding glomeruli per odor.

Another possibility is that OMP may be directly involved in axon guidance in OSNs. This hypothesis is prompted by the presence of OMP in OSN axons, where it might regulate the level of cAMP and therefore affect axon path finding. The OB glomerular map is dependent on varying levels of cAMP controlled by a non-olfactory G protein (Gs) in the OSN axons^33,34^ and adenylyl cyclase III^45,46^. It is thought that the activity profile of each OR can modulate the cAMP levels locally via Gs and in turn cAMP levels can affect expression of axon guidance molecules like neuropilin 1 (Nrp1)^34,47^ The cAMP level dependent regulation of glomerular positioning appears to be most relevant for global A-P positioning^48,49^

An additional step of local segregation of axons is thought to occur by different mechanisms that may rely on cell adhesion molecules^50–52^ For example, the level of neuronal activity determines the degree of expression of the homophilic adhesive proteins Kirrel2/Kirrel3^52^, which might aid local coalescence of homotypic axons. If activity levels are perturbed, then expression patterns of such proteins may be altered, leading to inadequate segregation. Since the level of cAMP signaling determines the expression of these “Type II molecules”, OMP could be involved in local axon guidance in the OB by regulating cAMP levels. This hypothesis is further corroborated by recent work connecting OMP to cAMP levels. Elevating intracellular cAMP levels by adding phospho-diesterase inhibitors restored normal response kinetics in OMP−/− mice^30^. This implies that OMP acts upstream of cAMP production in the OSN signal transduction pathway and that loss of OMP decreases cAMP levels, perturbing the normal expression of “Type II” molecules and, thus, preventing proper glomerular convergence. Future work will further explore how downstream mitral and tufted cells, which connect to a single glomerulus and relay information to downstream targets primarily regarding activity at this glomerulus, are affected by OSN glomerular heterogeneity.

### Conclusions & Future Directions

Our findings demonstrate that OMP is necessary for efficient segregation of axons in their target glomeruli and fulfillment of the one receptor-one glomerulus rule. Functionally, OMP^−/−^ mice presented glomeruli populated by microdomains of differing odor sensitivities, whereas in OMP^+/−^ mice, odor responses within a glomerulus tend to be homogeneous. Using gene targeted mice, we found that OMP^−/−^ mice exhibited both anatomical microdomains within glomeruli and duplicate glomeruli, and these phenotypes are infrequent in OMP^+/−^ mice. Our results indicate that OMP plays an important function in axonal refinement of the adult OB glomerular map.

## Methods

### Functional Preparation

Fourteen OMP^−/−^ and 8 OMP^+/−^ mice were anesthetized using a cocktail of ketamine/xylazine (90mg/kg+9mg/kg), supplemented every 45 minutes (30mg/kg+3mg/kg), and their heads fixed to a thin metal plate with acrylic glue. The dorsal aspect of the olfactory bulb was exposed through a small cranial window covered with 1.2% low melting point agarose and a glass coverslip to keep the tissue moist and to minimize motion artifacts. Heartbeat, respiratory rate, and lack of pain reflexes were monitored throughout the experiment. All animal procedures conformed to NIH guidelines and were approved by Harvard University’s Animal Care and Use Committee.

### Odorant Stimulation

A custom odor delivery machine was built to deliver up to 100 stimuli automatically and in any desired sequence under computer control of solenoid valves (AL4124 24 VDC, Industrial Automation Components). Pure chemicals and mixtures were obtained from Sigma and International Flavors and Fragrances. Odorants were diluted 100-fold into mineral oil and placed in blood collection tubes (Vacutainer, #366431) loaded on a custom made rack and sealed with a perforated rubber septum circumscribing two blunt end needles (Mcmaster, #75165A754). Fresh air was pumped into each tube via one needle by opening the corresponding solenoid valve. The mixed odor stream exited the tube through the other needle and was delivered at ~0.5 l/min via Teflon-coated tubing to the animal’s snout. For each stimulus, 20 seconds of odorant presentation were preceded and followed by 20 seconds of fresh air. At least 30 to 60 seconds of additional fresh air between stimuli was allowed. Each stimulus was delivered 1-3 times. A list of the odorants used in our widefield (98) and two photon (32) experiments is provided in Supplementary Table 1.

### Multiphoton Imaging

A custom-built two-photon microscope was used for *in vivo* imaging. SpH signals were imaged with a water immersion objective (20X, 0.95 NA, Olympus) at 910 nm using a Mira 900 Ti:Sapphire laser (8 W Verdi pump laser) with a 150 fs pulse width and 76 MHz repetition rate. The shortest possible optical path was used to bring the laser onto a galvanometric mirror scanning system (6215HB, Cambridge Technologies). The scanning system projected the incident laser beam through a scan lens and tube lens to backfill the aperture of an Olympus 20X, 0.95 NA objective. A Hamamatsu R3896 photomultiplier was used as photo-detector and a Pockels cell (350-80 BK and 302RM driver, Con Optics) as beam power modulator. The current output of the PMT was transformed to voltage, amplified (SR570, Stanford Instruments) and digitized using a data acquisition board that also controlled the scanning (PCI 6110, National Instruments). Image acquisition and scanning (2-5Hz) were controlled by custom-written software in Labview.

### Signal detection and analysis

Odor responses were calculated as average change (across 3-5 repeats) in fluorescence intensity divided by baseline (pre-odor) intensity using software written in IgorPro. ROIs were manually selected based on anatomy and odor responses. Care was taken to avoid selecting ROIs with overlapping neuropil. To facilitate detection of responding glomeruli, we calculated a ratio image for each odor (average of images in odor period minus average of images in pre-stimulus period, normalized by the pre-stimulus average). We further obtained a maximum pixel projection of all odor responses, assigning to each pixel in the field of view the maximum response amplitude across the odorants sampled – this allowed us to visually identify odor responsive regions. These responsive regions of interest mapped to individual glomeruli in the fluorescence image and were selected for further analysis.

We used principal component analysis (PCA) to identify pixels within the glomerulus that have similar response dynamics. Pixels above the signal threshold (2 SDs from baseline fluctuations) were projected on the plane determined by two of the first three principal components revealing functional clusters. To determine the spatial location of these clusters, we correlated the average response spectrum of each identified functional cluster to the odor response spectra of each pixel in the field of view. In a subset of experiments, the same analysis was extended to larger field of views (FOVs), across FOVs obtained from the same animal, as well as across animals within similar locations on the bulb surface.

### Gene-Targeted Mouse Lines

The generation of OMP knockout (OMP^−/−^) mice has been previously described in the literature, in which the OMP alleles have been replaced with the green synaptic fluorescent reporter SpH^35^. Generation of the *P2-IRES-taulacZ* and *M72-IRES-tauRFP_2_* mice using homologous recombination has also been described^2^.

OMP heterozygous (OMP^+/−^) mice were generated by breeding OMP^−/−^ mice with wild-type C57BL/6 mice (Jackson Labs) *P2-IRES-taulacZ* mice were bred with OMP^−/−^ mice to generate P2-lacZ mice that were heterozygous or homozygous for the OMP deletion. Similarly, *M72-IRES-tauRFP_2_* were bred with OMP^−/−^ mice to generate M72-RFP mice that were heterozygous or homozygous for the OMP gene deletion.

Genotyping for *P2-IRES-taulacZ, M72-IRES-tauRFP_2_*, and OMP SpH was carried out using DNA purified from toe/tail samples. PCR was conducted using primers in the Jackson Laboratories Mice Database and EconoTaq PLUS GREEN 2X Master Mix.

All mice were housed under a light/dark cycle of 12/12 hours and in accordance with institutional requirements for animal use and care.

### Whole-Mount Preparation

OMP^−/−^ or OMP^+/−^ whole mount brain was prepared by euthanizing the animal, dissecting out the skull, and removing bone to expose the dorsal surface of the OBs.

P2-lacZ/ OMP^−/−^ or OMP^+/−^ whole mount brain was prepared by first euthanizing the animal, dissecting out the brain from the skull, fixing the brain in 4% paraformaldehyde (PFA) in 1X phosphate-buffered saline (PBS) for 30-60 minutes on ice, and rinsing the brain in 1X PBS. The activity of the enzymatic reporter β-galactosidase (β-gal) was revealed by incubating the brain at 37° C overnight in a 1X PBS solution containing 5 mM potassium-ferricyanide, 5 mM potassium-ferrocyanide, and 1 mg/mL X-gal.

For M72-RFP mice, in both OMP^−/−^ and OMP^+/−^ backgrounds, whole mount brains were prepared by first euthanizing the animal and removing bone to expose the dorsal surface of the OBs. These brains were fixed in 4% PFA in 1X PBS for 30-60 min. on ice and then rinsed in 1X PBS.

### Sectioning

OMP-spH brains were fixed in 4% PFA in 1X PBS overnight and serial coronal sections (50 μm) of the OBs were taken using a Leica VT1000S microtome. Free-floating sections were mounted on VWR Superfrost Plus Micro Slides with VECTASHIELD Mounting Medium, which were then coverslipped and sealed with nail polish.

P2-lacZ and M72-RFP brains were processed for cryo-sectioning. Fresh (unfixed) brains were dissected out of the skull and transferred to a 30% sucrose solution in distilled water at 4 °C until they sank to the bottom of the container. Brains were then embedded in Tissue-Tek O.C.T. in a cryo-mold and rapidly frozen on dry ice. Samples were stored at −80 °C and placed in cryostat chamber at −16 °C for 1 hour prior to sectioning to achieve temperature equilibration. Serial coronal sections (16 μm) of the OBs were collected onto VWR Superfrost Plus Micro Slides. VECTASHIELD Mounting Medium with DAPI was placed on M72-RFP slides, which were then covered with a glass coverslip and sealed with nail polish. Immunohistochemistry, as described below, was performed on P2-lacZ sections before slides were covered with a glass coverslip with VECTASHIELD Mounting Medium with DAPI and sealed with nail polish.

### Immunohistochemistry

Following cryo-sectioning, immunohistochemistry was performed on P2-lacZ sections to visualize β-gal signal. Sections were equilibrated in 1X PBS for 5 min. at room temperature (RT), fixed in 4% PFA in 1X PBS for 10 min. at RT, and washed in 1X PBS for 2 × 3 minutes at RT. Sections were then incubated in 0.5% Triton-X-100 in 1X PBS for 30 minutes at RT, washed in 1X PBS for 3 × 5 minutes at RT, and incubated in 5% normal goat serum in 1X PBS (blocking buffer) for 30 minutes at RT. Finally, sections were incubated in primary antibodies in blocking buffer overnight at 4°C, washed in 1X PBS for 3 × 3 minutes at RT, and incubated in secondary antibodies in blocking buffer for 2 hours at RT.

The primary antibody used was rabbit anti-β-gal (Molecular Probes) at a dilution of 1:500 in blocking buffer. The secondary antibody used was goat anti-rabbit conjugated Alexa Fluor 594 (Life Technologies) at a dilution of 1:500 in blocking buffer.

### Imaging

All whole mount samples (OMP-spH, P2-lacZ, and M72-RFP) were imaged using the Zeiss Axio Zoom.V16, with varying exposure and magnification settings across samples. GFP fluorescence was monitored for OMP-spH whole mounts, reflected brightfield for P2-lacZ whole mounts, and RFP fluorescence for M72-RFP whole mounts.

Serial coronal sections (16μm) of P2-lacZ and M72-RFP OBs were imaged using the Zeiss Axio Scan.Z1. For P2-lacZ sections, fluorescent channels included GFP, DAPI, and Alexa Fluor 594, with exposure times of 200 ms, 5 ms, and 50 ms, respectively. For M72-RFP sections, fluorescent channels included GFP, DAPI, and RFP, with exposure times of 200 ms, 5 ms, and 100 ms, respectively. Tiled images of each section were generated at a magnification of 10x for both P2-lacZ and M72-RFP sections. Images were not modified other than to balance and adjust brightness and contrast.

### Heterogeneity Overlap and Glomerular Size Analysis

For the quantification of heterogeneity within a single glomerulus, custom software (MATLAB) was created to analyze the overlap between red (β-gal or RFP) signal labeling single odorant receptor identified glomeruli and green (OMP-spH) signal, which labeled all glomeruli. A threshold based on background pixels (selected outside of glomerular regions) was determined, highlighting glomeruli. Glomerular regions of interest were selected in which the number of overlapping red/green pixels, only green pixels, and background pixels was determined. From these pixel measurements, glomerular homogeneity could be expressed as a fraction of the number of overlapping red/green pixels divided by the number of all spH positive pixels in a single glomerular structure. For the glomerular size analysis, custom software (MATLAB) was created to calculate the area in pixels within a selected glomerular region of interest. These glomerular areas in pixels could be readily expressed in μm^2^ using a pixel-to-μm conversion factor.

---

**Supplemental Figure 1.**
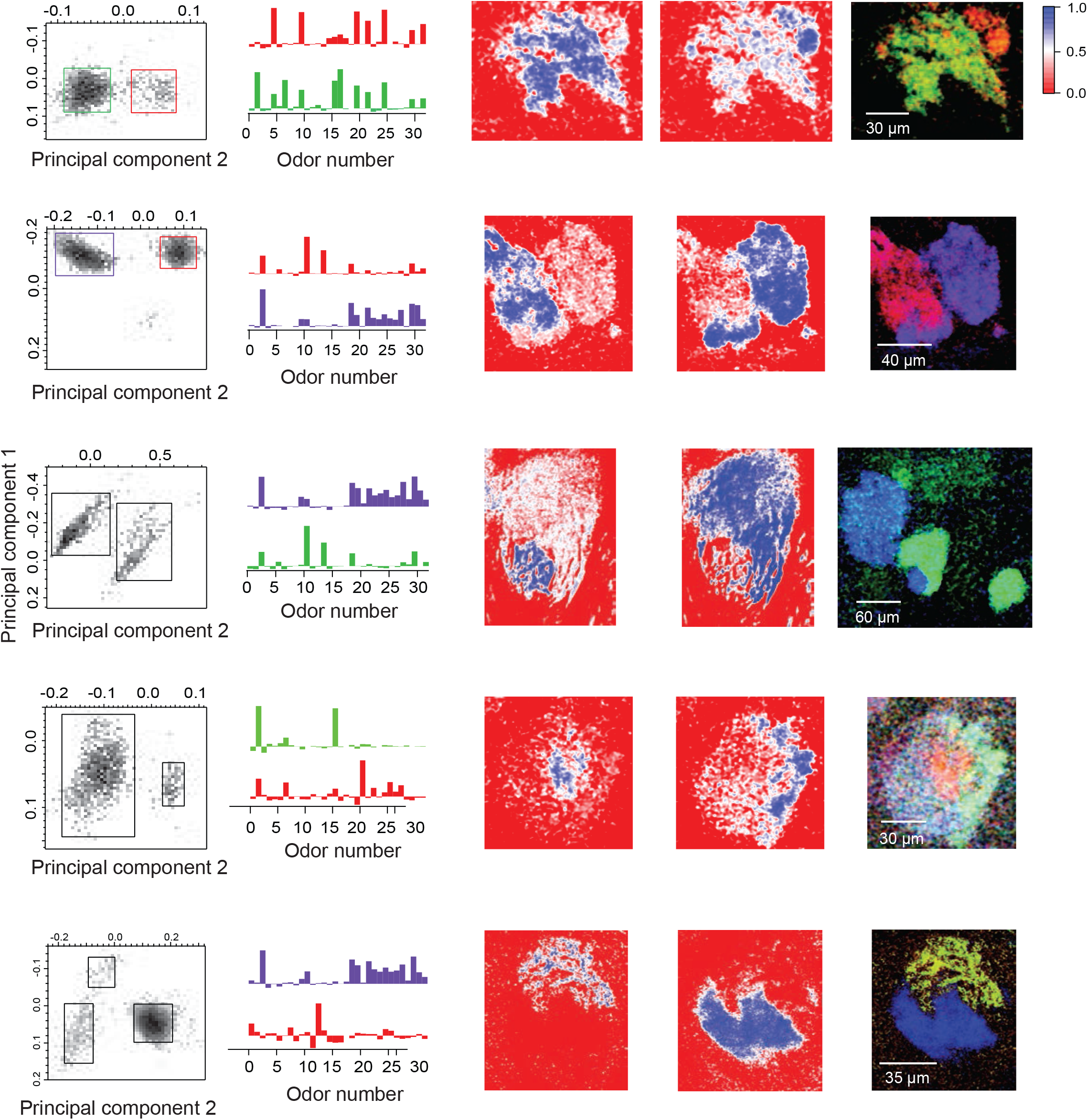
Examples of functional glomerular heterogeneity in OMP^−/−^ mice. (*Left*) Projection of all pixels in each of the example fields of view on two principal components (PC1 vs. PC2) coordinates. PCA was performed on the odor response spectra of each pixel. (*Center*) Average odor response spectra of the functional clusters identified by PCA. (*Right*) Correlograms showing the spatial distribution of the pixel clusters identified via PCA within the example glomeruli. These were obtained by correlating the average odor response vectors of the identified clusters to the odor responses of individual pixels in each field of view. Note that each functional cluster corresponds to a spatially contiguous area, filling either a spatial microdomain within a larger heterogeneous glomerulus, or occupying the whole anatomical glomerulus. RGB color scheme overlays the correlograms corresponding to the sub-glomerular clusters.

**Supplemental Figure 2.**
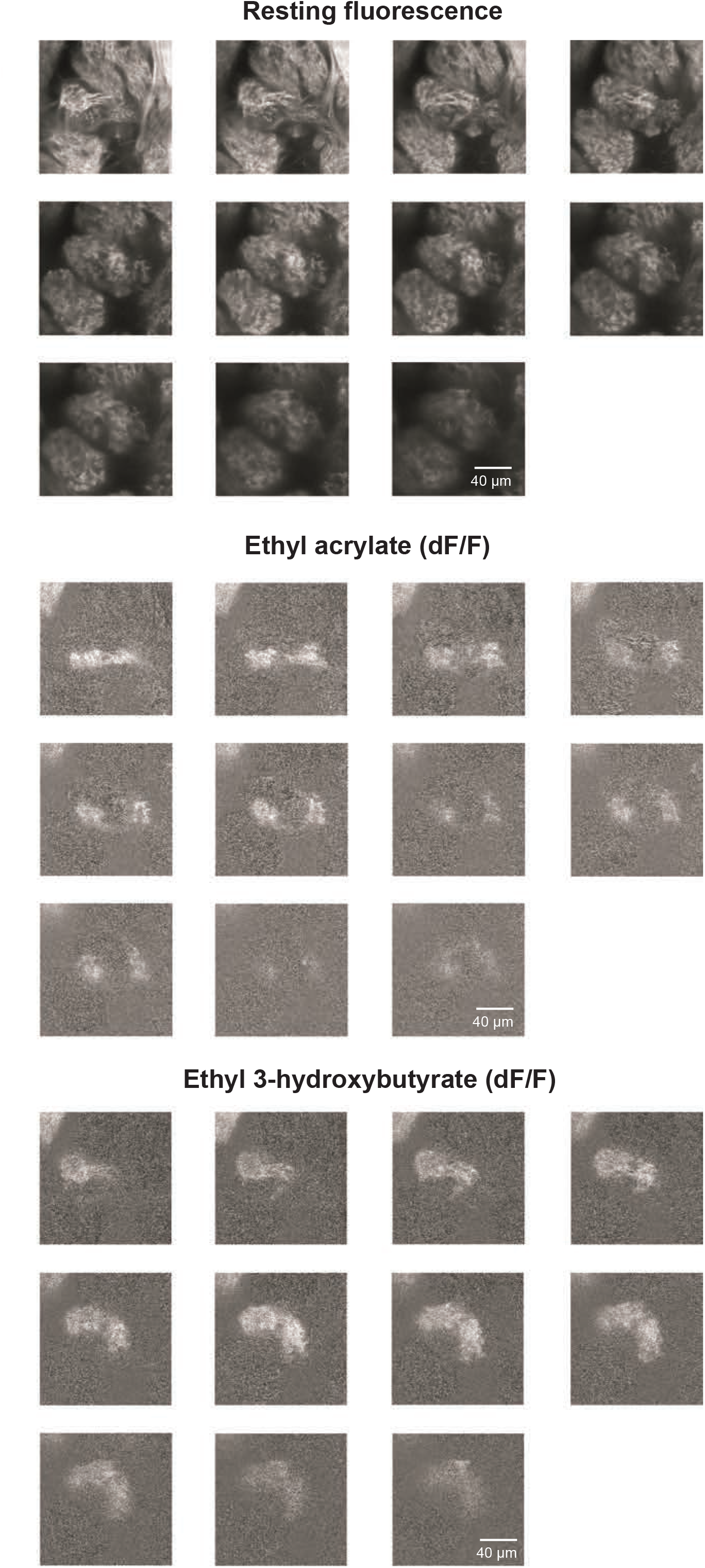
Z-stack through a functionally heterogeneous glomerulus. (*Top*) Resting fluorescence of OMP-spH labeled glomeruli sampled 8μm apart along the z-axis. Note the different resting fluorescence intensity in different regions of the glomerulus. (*Center*) dF/F ratio images in response to ethyl acrylate delivery indicate two functional microdomains across different z-planes throughout the glomerulus. (*Bottom*) dF/F responses to ethyl 3-hydroxybutyrate (E3HB). A distinct microdomain from the one shown above (*Center*) was activated by E3HB.

**Supplemental Figure 3.**
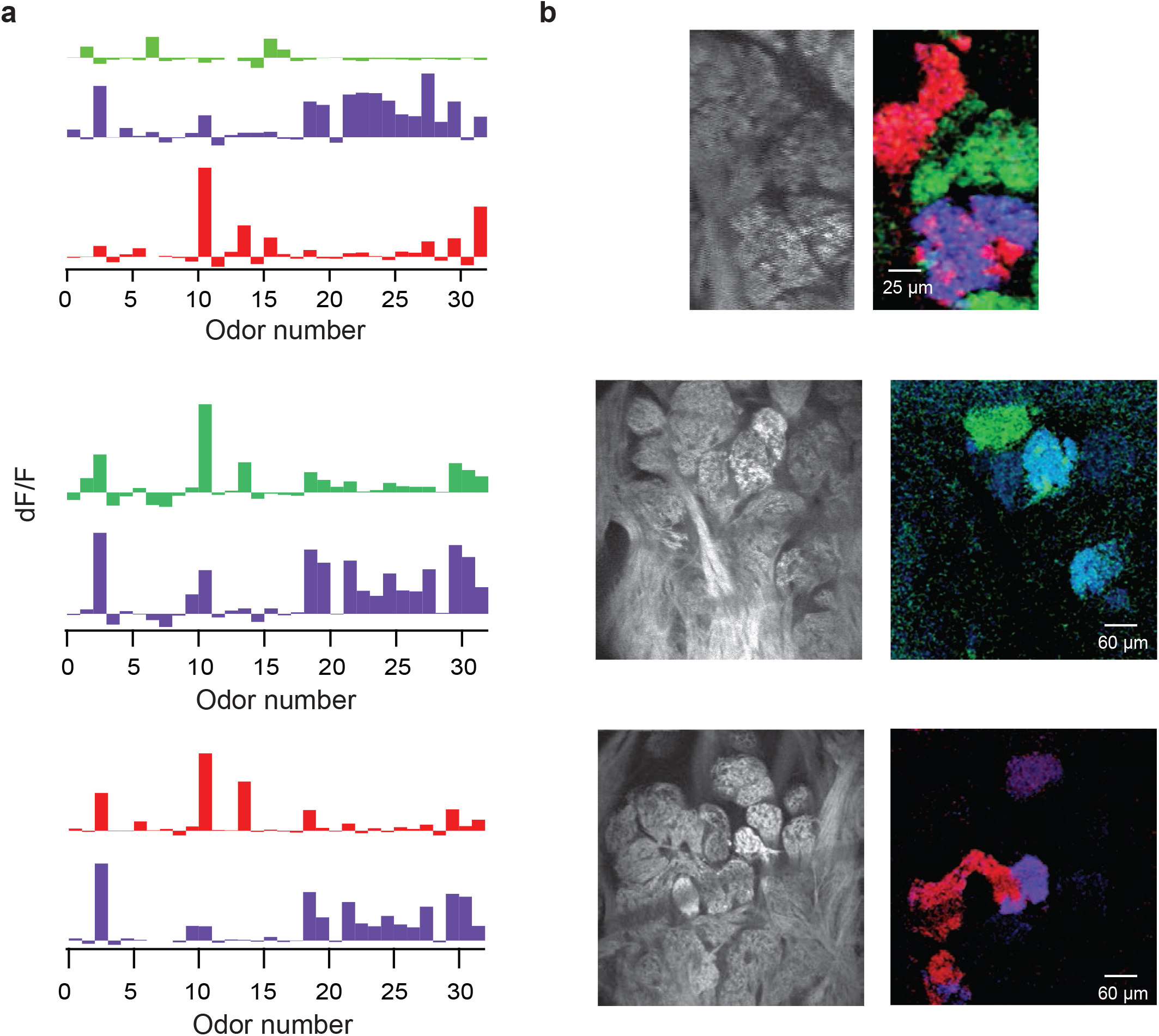
Local functional glomerular duplications in OMP^−/−^ mice. Three example fields of view revealing functionally heterogeneous glomeruli and local duplicates. (*Left*) Example average odor response spectra of PCA identified functional clusters corresponding to glomerular microdomains. (*Right*) Resting fluorescence and overlay of correlograms. Note the presence of nearby functionally homogeneous and heterogeneous glomeruli. For example, in the top panel are depicted five anatomically identifiable glomeruli. The correlograms (*Right*) indicate that red coded microdomains are shared between two close-by anatomical glomeruli. Blue and red microdomains mix within one anatomical glomerulus. The green clusters are present as two homogeneous nearby glomeruli.

**Supplemental Figure 4.**
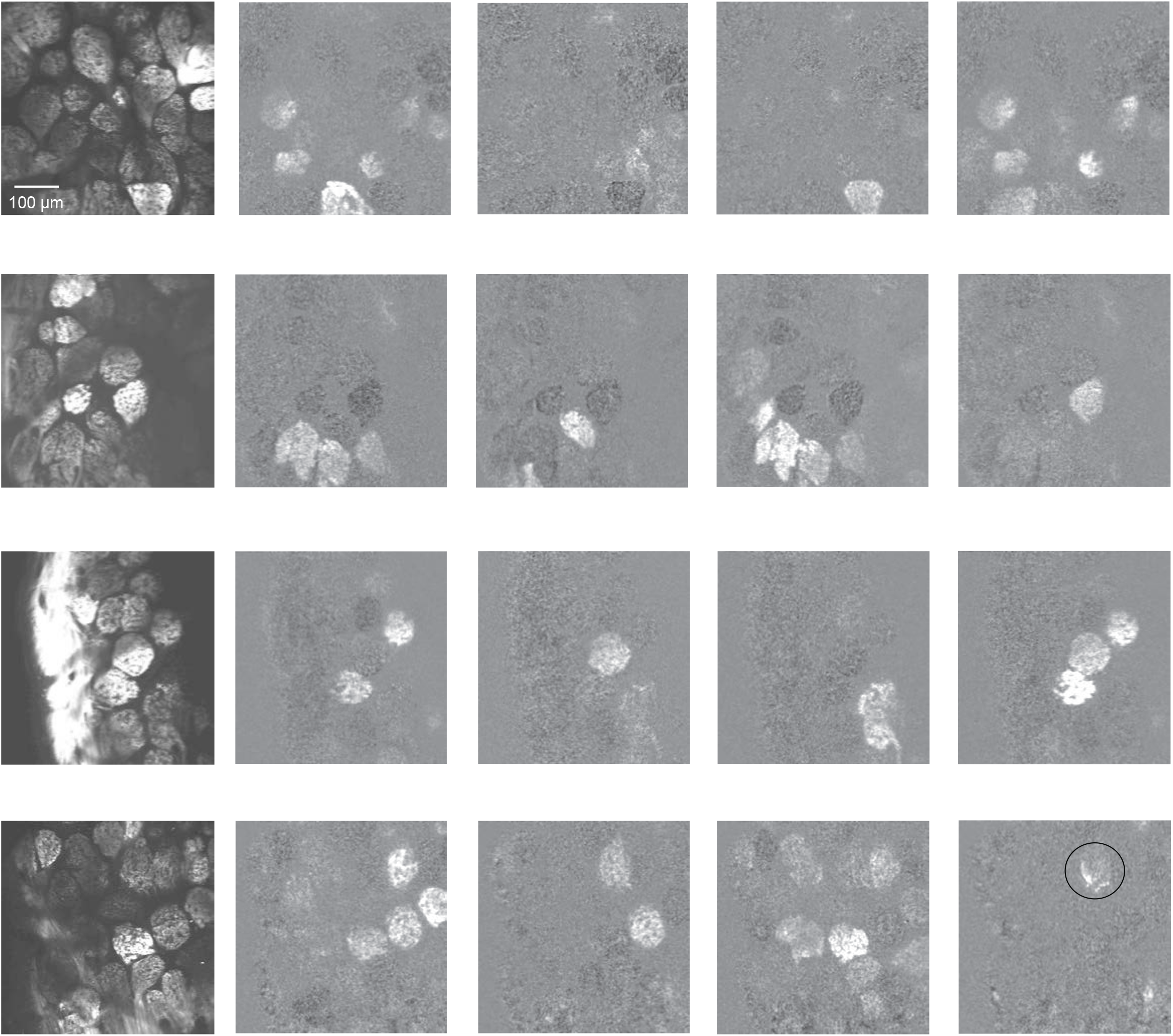
Examples of glomerular responses in OMP^+/−^ mice. Four example fields of view from three olfactory bulb hemispheres. (*Left*) Resting fluorescence of spH labeled glomeruli in OMP^+/−^ mice. (*Right*) Example glomerular odor responses (dF/F) to a panel of 32 stimuli from the fields of view shown on the left. Circle marks putative heterogeneous glomerulus.

**Movies 1 & 2**.

Z-stacks of resting fluorescence through two example functionally heterogeneous glomeruli.

**Movies 3 & 4**.

Example dF/F responses to ‘fresh air’ and different odorants in the panel within one optical plane chosen within the 2 glomeruli shown in Movies 1 & 2.

**Movies 5,6 & 7**.

Example functionally heterogeneous glomerulus. Z-stacks of resting fluorescence and dF/F responses to 2 odorants in the panel (ethyl acrylate and ethyl 3-hydroxybutryate) across 38 optical planes sampled 2μm apart along the z-axis with respect to the bulb surface.

